# Mechanical stretch disrupts calcium dynamics and redistributes Piezo1 in human astrocytes

**DOI:** 10.1101/2025.10.08.681038

**Authors:** Shahrzad Shiravi, Akash Chakka, Xi Xiao, Meilin Fernandez Garcia, Alexandra Yufa, Angela Mitevska, Carina Seah, Laura M. Huckins, Kristen J. Brennand, John D. Finan

**Affiliations:** Department of Mechanical and Industrial Engineering, University of Illinois Chicago, Chicago, IL 60607, USA; Department of Psychiatry, Yale University School of Medicine, 34 Park Street, New Haven, CT 06520, USA; Department of Biomedical Engineering, University of Illinois Chicago, Chicago, IL 60607, USA; Pamela Sklar Division of Psychiatric Genomics, Departments of Psychiatry and of Genetics and Genomic Sciences, Icahn School of Medicine at Mount Sinai, New York, New York, 10029 USA

**Keywords:** Traumatic brain injury, hiPSC-derived astrocytes, calcium dynamics, mitochondrial dysfunction, Piezo1, RNA sequencing

## Abstract

Astrocytes regulate the activity of nearby neurons so disruption of astrocyte calcium dynamics by traumatic brain injury (TBI) could have profound consequences for neural network activity in the brain. In this study, human induced pluripotent stem cell (hiPSC)-derived astrocytes were used in a two-dimensional (2D) *in vitro* stretch injury model to evaluate the effect of trauma on calcium dynamics, mitochondrial function, and the mechanosensitive ion channel Piezo1. Outcomes were assessed using live imaging, immunostaining, and RNA sequencing. Cell viability, mitochondrial membrane potential, and spontaneous calcium transients declined as injury severity increased. At moderate injury severity, the decreases in mitochondrial membrane potential and calcium dynamics were temporary. The spatial distribution of Piezo1 also changed temporarily after injury. RNA sequencing identified 196 genes that changed expression after injury, including downregulation of mitochondrial and oxidative metabolic processes and upregulation of cortical thinning pathways. These findings establish this model as a platform for investigating the cellular mechanisms of TBI and its influence on neurodegeneration.

## Introduction

Traumatic brain injury (TBI) is a global public health concern, affecting around 69 million individuals annually[1] and contributing to long-term neurological impairments[2]. Mild TBI, which accounts for the majority of cases[1], has been linked to long-term consequences, including increased risk of psychiatric disorders[3] and neurodegenerative disorders such as Alzheimer’s disease[4]. The biological cascade triggered by TBI includes not only immediate structural damage, but also delayed and diffuse processes such as calcium dysregulation, mitochondrial dysfunction, and cytoskeletal remodeling [5], [6]—many of which may evolve over time and contribute to chronic pathology [7], [8].

Clinical functional magnetic resonance imaging (fMRI) studies have provided invaluable insight into the functional consequences of TBI but face inherent limitations in capturing fast, cell-specific changes in signaling events [9]. Animal models are constrained by species-specific brain structures, cell composition, and injury responses [10]. Human induced pluripotent stem cell (hiPSC)-derived astrocytes reproduce human pathophysiology and permit standardization and manipulation of genetic factors affecting outcome [11], [12], [13]. Two-dimensional stretch-based models are well-suited for isolating the effects of mechanical strain - a primary driver of TBI pathology [14] - on individual cell types [15]. The application of *in vitro* stretch injury to hiPSC-derived astrocytes permits full control of mechanical and genetic factors driving neurotrauma pathology in a human system.

Astrocytes play a central role in the brain, regulating extracellular ion balance, providing metabolic support, and mediating calcium signaling so they are tightly linked to neuronal stability and network activity [16], [17], [18], [19], [20]. In this study, we used a previously developed 2D stretch injury model [24], [25] to examine the structural and functional responses of mature hiPSC-derived astrocytes to a range of mechanical injury severities. Using live imaging and custom image analysis tools, we quantified mitochondrial membrane potential, cell viability, and calcium transients after trauma. Our results reveal strain-dependent alterations in mitochondrial function and calcium dynamics, as well as changes in the spatial organization of the cytoskeleton and Piezo1 channels. These findings underscore the sensitivity of astrocytes to mechanical input and highlight the value of human-based *in vitro* models for dissecting the cell-intrinsic responses that determine TBI morbidity.

## Methods

### Quantification of membrane stretch

Stretchable Plates were fabricated by bonding bottomless Nunc plate frames (Thermo Fisher Scientific, 12-565-600) to polydimethylsiloxane (PDMS) membranes (SMI Manufacturing, .010” NRV G/G 40D TT) using a previously described method [24], [25].The PDMS membranes were plasma treated, aligned to the plate frames, clamped, and cured for 24 hours. For quantifying membrane stretch, a plate was airbrushed with carbon black ink so that the deformation of the membrane could be visualized. Then, it was indented to various depths using the same settings that were used to injure cultures while the plate bottom was imaged at 1500 frames per second with a Photron FastCam Mini WX100 high speed camera. GOM Correlate image correlation software was used to quantify the nominal Green strain in these images. Strain histories were calculated for multiple points in each well and then their mean was taken to calculate a single strain history that was used to find the peak strain in that well.

### Cell culture

hiPSC-derived astrocytes were generated in Dr. Kristen Brennand’s laboratory at Yale University using a previously published method [26], [27]. Briefly, hiPSCs were differentiated into astrocytes using a lentiviral vector that rapidly induced Sox9 and NFIB transcription factors. Cells were maintained in StemFlex™ Basal Medium (Thermo Fisher Scientific, A3349301), DMEM/F12 with HEPES (Thermo Fisher Scientific, 11330032), and FGF-enriched medium containing Neurobasal™ medium (Thermo Fisher Scientific, 21103049), B27 supplement (17504044), and CNTF (PeproTech, 450-13) for 7 days. At this stage, cells were dissociated using Accutase (Innovative Cell Technologies, AT104), cryopreserved in freezing medium (FGF medium with 20% DMSO), and shipped to Finan Lab at University of Illinois Chicago where they were thawed and seeded on custom 96-well PDMS plates. PDMS plates with sterilized by 15-minute immersion in 70% ethanol, followed by a 30-minute rinse with sterile deionized water, and coated with Geltrex. Plates were incubated overnight at 37 °C in a humidified incubator with 5% CO_2_. The following day, cryovials of cells were warmed in a 37 °C bead bath for 5 minutes, transferred to 4 mL of DMEM (Thermo Fisher Scientific,11965092), centrifuged at 400 × g for 5 minutes, resuspended in FGF medium ( Neurobasal™ medium, 2% B27 supplement (12587010), 1% GlutaMAX™ (Thermo Fisher Scientific, 35050061), 1% MEM Non-Essential Amino Acids (Thermo Fisher Scientific, 11140050), 1% sodium pyruvate (Thermo Fisher Scientific, 11360070), 8 ng/mL FGF2 (PeproTech, 100-18B), 5 ng/mL CNTF, 10 ng/mL BMP4 (PeproTech, 120-05ET), and 2.5 µg/mL doxycycline (Thermo Fisher Scientific, D9891)) and seeded at a density of 15k cells/well. Beginning on day 8, cultures were gradually transitioned to a maturation medium consisting of a 1:1 mixture of DMEM/F12 (Thermo Fisher Scientific,11320033) and Neurobasal™ medium, supplemented with 1% GlutaMAX™, 1% N-2 supplement (17502048), 1% sodium pyruvate, 5 µg/mL N-acetyl-cysteine (Sigma-Aldrich, A8199), 5 ng/mL heparin-binding EGF-like growth factor (Sigma Aldrich, E4643), 10 ng/mL CNTF, 10 ng/mL BMP4, 500 µg/mL dibutyryl cyclic-AMP (Sigma-Aldrich, D0627), and 2.5 µg/mL doxycycline. From day 10 onward, half-medium changes with maturation media were performed every 2–3 days. The hiPSC line used in this study was validated by flow cytometry for SSEA-4 and TRA-1-60 and by immunofluorescent imaging of SOX2, NANOG, OCT4, and TRA-1-60. WiCell Cytogenetic Services performed G-banded karyotyping to rule out chromosomal abnormalities. The MycoAlert kit (Lonza) was used routinely (every 2–4 weeks) to confirm that the cells were not contaminated with mycoplasma. All 2D stretch injuries were conducted after day 21 of culture, when astrocytes were considered mature and showed calcium transients (Supplementary video 1). Cells were immunostained for S100B and GFAP at day 21 to confirm astrocyte identity (Fig. S1).

### Stretch injury experiments

Mature astrocytes were subjected to mechanical injury using a previously developed custom-designed 2D injury machine capable of delivering controlled deformation in a 30ms period [25], [28]. The device uses lubricated, Teflon-coated, aluminum cylindrical posts to indent and stretch a flexible PDMS membrane upon which cells are cultured. The 30 ms timescale was selected to simulate the rapid mechanical loading characteristic of TBI because the rate of deformation influences the severity of the pathology [29]. To measure the effects of injury severity, five indentation depths were used: sham (no injury), 0.5 mm, 1.5 mm, 2.5 mm, and 3.2 mm. These depths were selected according to previously characterized strain values,[24], [25], [30], [31] to induce a range of mechanical strains that would cause injuries ranging from mild to severe in the cell monolayer. In the sham condition, the posts contacted the membrane without deforming it. The contact position was found by monitoring the well bottom with an overhead camera for evidence of contact as the posts approached. Following injury, cultures were returned to standard incubation conditions at 37 °C with 5% CO_2_ for 24 hours prior to further analysis.

### Live staining and imaging

Stains used in this study included Calbryte 520 AM (5 μM, AAT Bioquest, 20653) to visualize calcium dynamics, Hoechst (1:500 dilution, Thermo Fisher Scientific, H21486) to stain nuclei, tetramethylrhodamine(TMRM, 0.1 μM,Millipore Sigma, T5428) to evaluate mitochondrial membrane potential, and calcein AM (2 μM, Thermo Fisher Scientific, C1430) to evaluate cell viability. After injury, cells were incubated with the appropriate staining solution for 1 hour at 37 °C with 5% CO_2_ and washed three times with DPBS prior to imaging with a Olympus FV3000 confocal microscope. To measure live calcium dynamics, time series images were acquired from the Calbryte-stained wells, capturing 277 frames at a frequency of 1 Hz.

### Immunostaining

For all immunostaining experiments, cells were washed with DPBS and fixed using 4% paraformaldehyde (Avantor ScienceCentral, AAJ61899-AK) for 10 minutes at room temperature. After removing the fixative, cells were incubated with a blocking solution (5% normal donkey serum (Jackson ImmunoResearch Laboratories Inc., 017-000-121c), 1% bovine serum albumin Fraction V (Thermo Fisher Scientific, 15260037), and 0.2% Triton X-100 (Millipore Sigma, X100-5ML) in DPBS) for 30 minutes at room temperature. Following blocking, cells were incubated overnight at 4 °C with primary antibodies diluted in blocking solution. The primary antibodies used in this study were Piezo1 (monoclonal, 1:100; Thermo Fisher Scientific, MA5-32876), Anti-S100B ( 1:100, Sigma Aldrich, S2532) and Chicken Anti-GFAP (1:100, Aveslabs). The next day, cells were washed twice with cold DPBS and incubated for 1 hour at room temperature with secondary antibodies diluted in blocking solution. The secondary antibodies included Alexa Fluor 647 Donkey Anti-Mouse secondary (1:200; Jackson, 715-605-151) and Alexa Fluor 488 Donkey Anti-Chicken ( 1:200, Jackson, 703-545-155). Phalloidin (165 nM; Thermo Fisher Scientific, A12379) was also used to stain actin fibers. DAPI (1:500; Millipore Sigma, 10236276001) was then applied for 5 minutes for nuclear counterstaining. Finally, cells were washed three times with DPBS. During these steps, cells were protected from light to prevent photobleaching.

### Image analysis

Image analysis was performed using CellProfiler 4.0 and MATLAB. Nuclear segmentation, essential for assessing cell viability and mitochondrial potential, was based on Hoechst staining. To exclude nuclear fragments and ensure accurate identification of intact nuclei, a two-step Otsu thresholding strategy was used. Intact nuclei were further filtered based on solidity to remove irregular or overlapping shapes. For cytoplasmic signal analysis (TMRM and Calcein AM), nuclei were expanded to define somatic regions of interest. At 7 days post-injury, TMRM images were acquired but could not be quantified due to a technical issue that resulted in loss of the DAPI channel, preventing accurate cell identification.

To quantify calcium activity across the entire field of view, a custom approach was developed using difference images - generated by subtracting each frame from the next (e.g., frame 2 - frame 1) - to highlight regions of increased fluorescence associated with calcium transients. These images were segmented using global minimum cross-entropy thresholding (Fig S2). Small artifacts were excluded using a size filter, and the area of each transient was quantified. The sum of these areas across all frame pairs was defined as the Asynchronous Activity Index (AAI), a metric reflecting the overall frequency and spatial extent of calcium activity in each recording. The AAI method was used to analyze recordings from wells that did not have nuclear staining.

In recordings that had nuclear staining, the mean fluorescence intensity of the Calbryte signal overlapping each nucleus was extracted in MATLAB. Calcium transients were identified using the pre-existing findpeaks function in MATLAB. Calcium response was quantified as the difference in peak count (Δ peak count) between the pre-treatment and post-treatment conditions.

### Pharmacological Activation of Piezo1

To assess Piezo1 presence and functionality in hiPSC-derived astrocyte cultures, cells were stained with 5 µM Calbryte 520 AM (AAT Bioquest, 20653) for one hour and treated with varying concentrations of Yoda1 (Millipore Sigma, SML1558-5MG), a small-molecule agonist that selectively activates the Piezo1 mechanosensitive calcium channel. Experimental conditions included sham, vehicle, and Yoda1 treatments at 1 µM, 5 µM, 10 µM, and 20 µM. The calcium-dependent fluorescence was recorded using confocal microscopy to evaluate Piezo1 activation. First, baseline activity was imaged at 1Hz for 150 frames (2.5 min). Imaging was then briefly paused (∼1 min) to allow addition of Yoda1 or vehicle. Then, imaging immediately resumed for another 150 frames to capture the response.

### RNA extraction

RNA samples were harvested using TRIzol-LS (Thermo Scientific, 10296028), and total RNA was isolated with the RNeasy Plus Micro Kit (Qiagen, 74104) according to the manufacturer’s protocol. RNA concentration and integrity were assessed using a NanoDrop 2000 spectrophotometer (Thermo Scientific) and an Agilent 2100 Bioanalyzer (Agilent), respectively.

### RNA-seq data generation

A low-input RNA-seq protocol was used to generate transcriptomic data from hiPSC-derived astrocytes. Poly(A)-enriched RNA was processed for library preparation using the SMART-Seq v4 Kit (Takara Bio). Libraries were sequenced using a paired-end 150 base pair configuration, with approximately 40 million reads generated per sample.

### RNA-seq data preprocessing

FASTQ files were processed using the procedure described by Ritchie et al. [32]. Short reads with Illumina adapters were trimmed and low quality reads were removed with TrimGalore [33]. The remaining high-quality reads were then aligned using the hg38 reference genome and the universal RNA-seq aligner STAR 2.7.9a with default parameters [34]. Bam files were organized with Samtools and quantified with featureCounts [35] with specific parameters (-T 5, -t exon and -g gene_id). Raw count data was converted to counts per million (CPM) in R using the cpm function in the edgeR [36] package to normalize for library size and sequencing depth. Genes with CPM values greater than 2 in 40% or more of samples were retained for analysis.

### Differential gene expression analysis

Transcriptional signatures were generated using methodology adapted from Hoffman et al. [37], using scripts available at www.synapse.org/hiPSC_COS. Gene expression values were normalized using Voom [32]. The voom-transformed data was visualized to assess normalization quality. For differential gene expression analysis of trauma/sham astrocytes, a moderated t-test was applied using limma [32] . To correct for multiple testing including multiple independent replicates per donor, the DREAM [38] function from the variancePartition package was used. Gene-level significance was adjusted for multiple comparisons using the Benjamini-Hochberg method to control the false discovery rate (FDR). Genes meeting FDR < 5% were considered significantly differentially expressed.

### Gene set enrichment analyses

Significant DEGs (pFDR<0.05) were partitioned into up-regulated and down-regulated genes. Gene set enrichment was performed separately in each group, testing for enrichment of genes in gene sets taken from (1) gene ontology biological processes (GOBP)[39], [40], (2) GWAS catalog [41].

Gene set enrichment was performed using the FGSEA R package [42], with the background set consisting of all genes retained after expression filtering. Benjamini– Hochberg false discovery rate (FDR) correction was applied independently within each gene set collection to correct for multiple testing.

## Results

### Membrane strain increased with increasing indentation depth

The actual indentation depth was within 6% of the prescribed indentation depth for all depths and was highly repeatable (see Table 1). The peak membrane strain values increased as the indentation depth increased. They were also quite repeatable, with standard deviation ranging from 6 to 12% of the mean. The injury process induced a homogeneous, equibiaxial strain field across the culture surface, ensuring that cells within the indentation area experienced uniform mechanical strain (see Fig. S4).

**Table 1:**
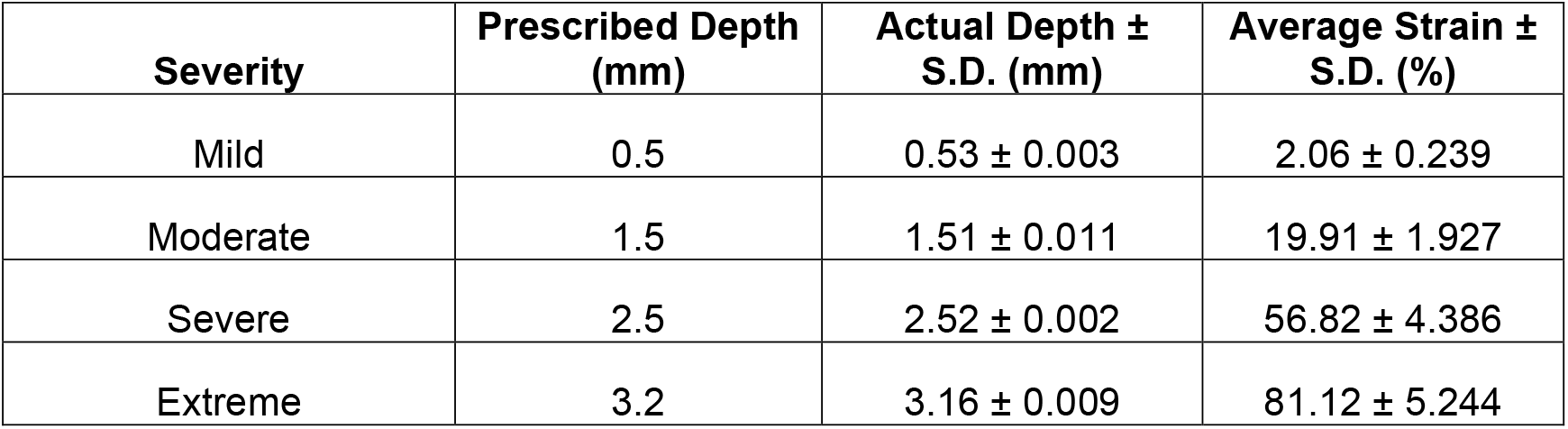
Indentation depths and associated peak membrane strain values (n = 21 wells, N = 3 plate stretch procedures).

### Cell viability, mitochondrial membrane potential, and calcium dynamics correlated with injury severity

Cell viability, mitochondrial membrane potential, and calcium dynamics declined progressively with increasing injury severity (Fig. 1). Mild injury did not alter cell viability or calcium dynamics but did cause a modest but statistically significant decline in mitochondrial membrane potential (p<0.5, Fig.1 C). Due to non-normal data distribution (confirmed using the Lilliefors test in MATLAB), statistical comparisons were performed using the nonparametric Kruskal-Wallis test [43], which revealed significant group differences (p < 0.05).

**Figure 1:**
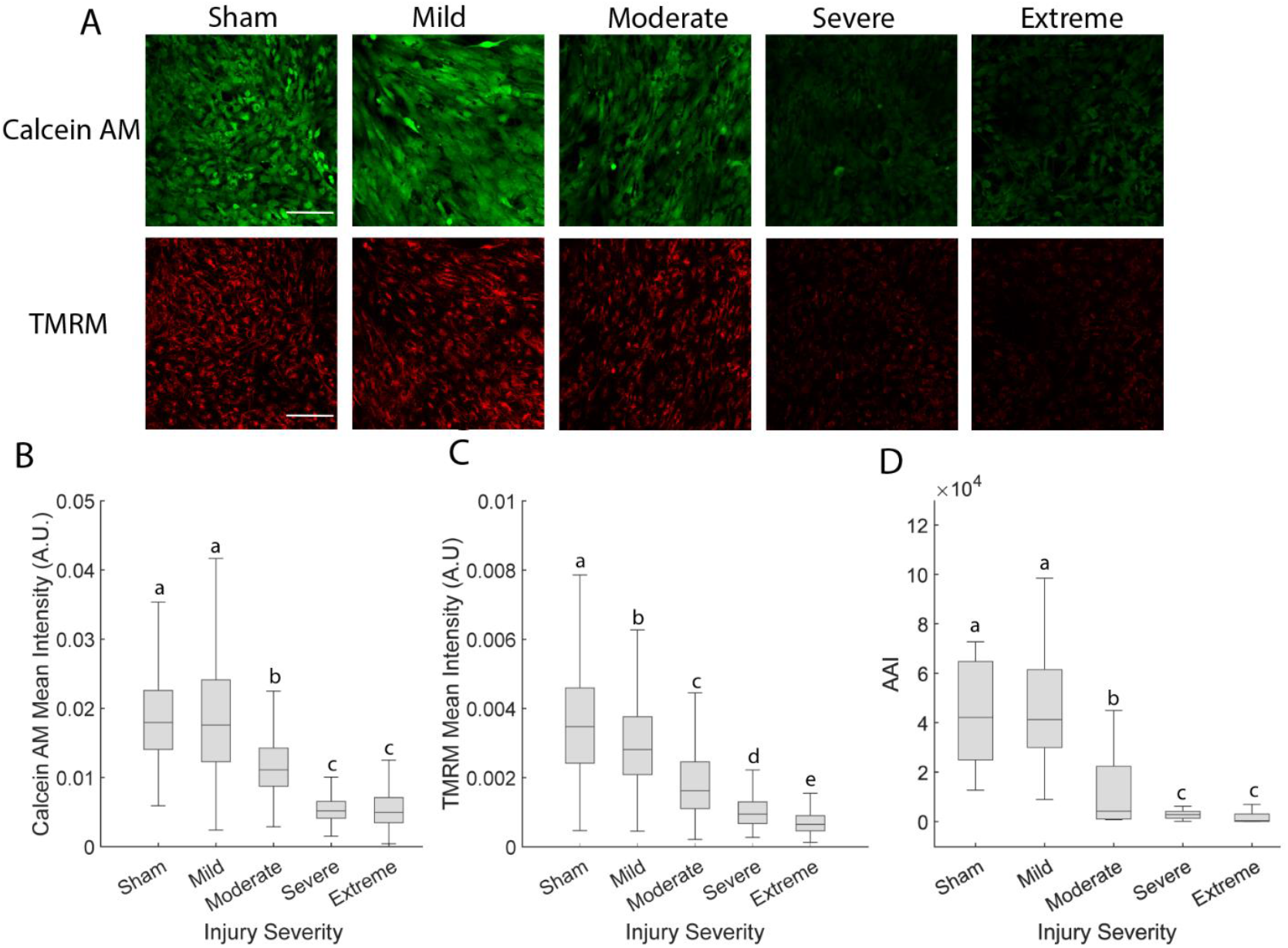
Cell viability, mitochondrial membrane potential, and calcium dynamics depend on trauma severity. (A) Fluorescent images of Calcein AM and TMRM staining (images are representative of 10 wells, scale bar = 200 μm). (B) Quantification of Calcein AM intensity (N=10 wells, n>=82 cells). (C) Quantification of TMRM intensity (N=10 wells, n>=82 cells). (D) Quantification of calcium dynamics (n=10 wells). Box plots display the median and interquartile range (25th–75th percentiles).Whiskers represent the range of values within 1.5 times the interquartile range. Groups that do not share a letter are significantly different (Kruskal-Wallis test, p< 0.05).

### Cultures made a partial recovery after Moderate injury

Following Moderate stretch injury, mitochondrial membrane potential increased at 1 hour, decreased significantly at 24 hours, and partially recovered by 72 hours without returning to sham levels (p<0.05, Fig. 2A, B). Calcium dynamics declined to zero by 24 hours post-injury but recovered to be indistinguishable from sham conditions by 7 days post-injury (Fig. 2C). Due to non-normal data distribution (Lilliefors test), a Kruskal-Wallis test was used and revealed a significant effect of time point (p < 0.05), with post hoc Bonferroni-corrected comparisons confirming group differences.

**Figure 2:**
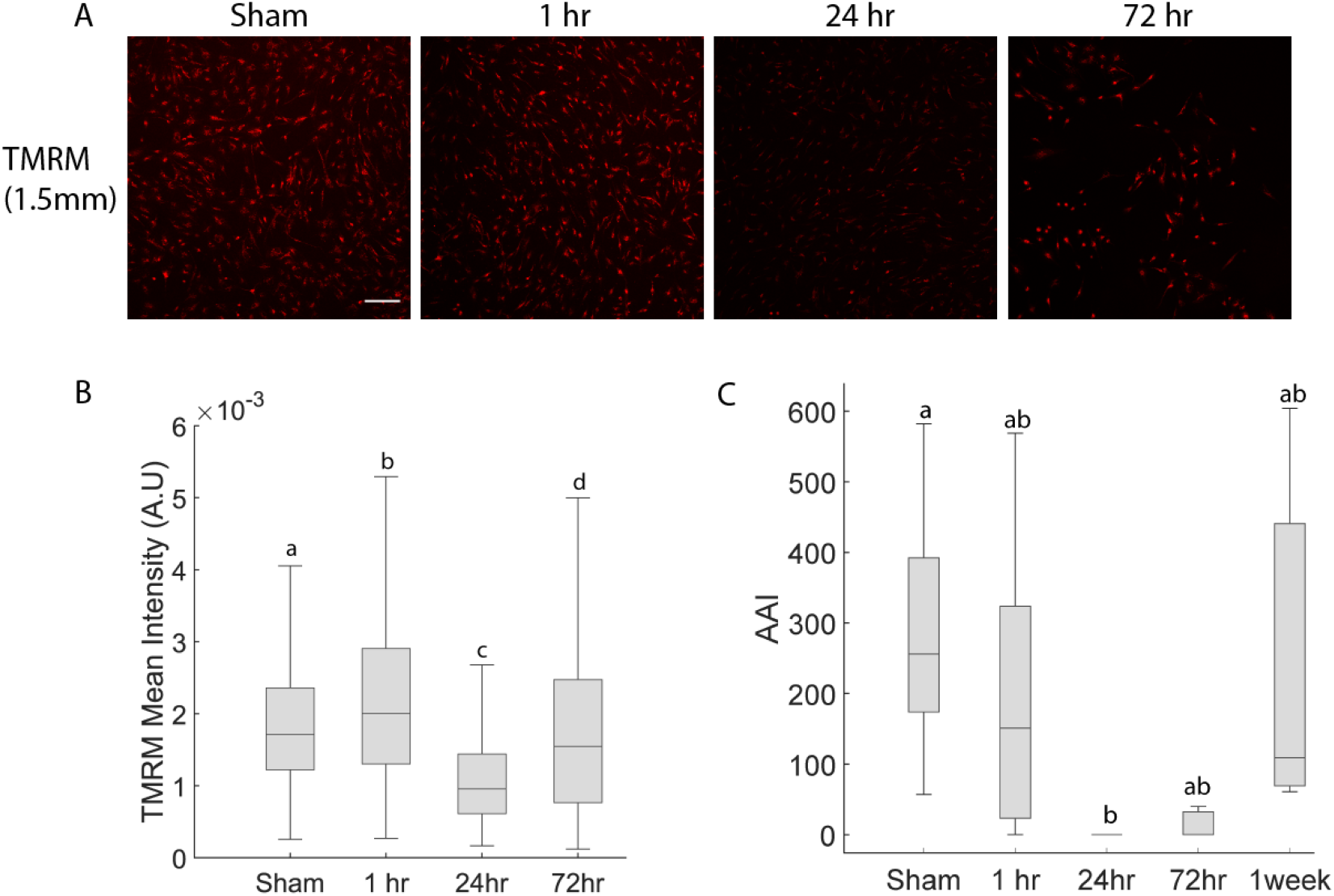
Temporal evolution of injury phenotype. (A) Representative TMRM images over a 1-week period post-injury (images are representative of 6 wells, scale bar = 100 μm). (B) Quantification of TMRM intensity over time (N=6 wells, n>=115 cells). (C) Quantification of calcium dynamics over time (n=6 wells). Box plots display the median and interquartile range (25th–75th percentiles).Whiskers represent the range of values within 1.5 times the interquartile range.

### Cell area declines after trauma

Since it was difficult to distinguish cells from their neighbors in monolayer cultures, cell area was evaluated by dividing the total area of phalloidin-stained cytoplasm by the number of nuclei in the field of view. Cell area declined modestly in the Mild and Moderate injury conditions 24 hours after injury of various severities (Fig. 3A, B). Surprisingly, it did not decline in the Extreme injury condition. However, it is worth noting that the overall cell count dropped dramatically in this condition (see Fig.S3) so this result may be influenced by a selection effect (i.e. only the most strongly attached cells survived 24 hours after Extreme injury and were included in the analysis). Cel area declined significantly 15 minutes after Moderate injury and recovered to sham level 72 hours after trauma (p<0.05, Fig. 3 C,D).

**Figure 3:**
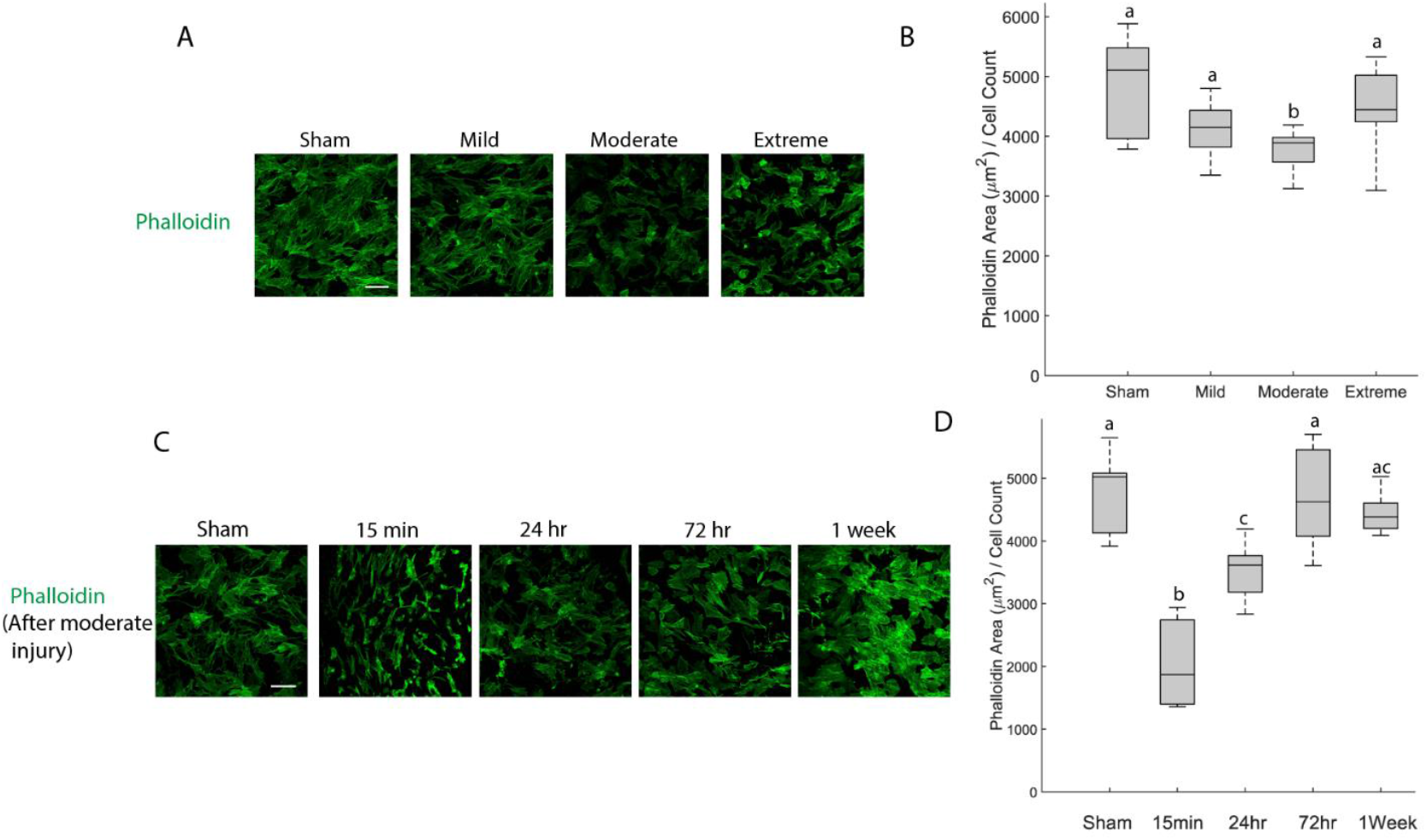
(A) Fluorescent images of phalloidin stained cultures 24 hours injury of various severities (images are representative of 6 wells, scale bar = 200 μm). (B) Total area of phalloidin-positive cytoplasm/nuclei count 24 hours injury of various severities. (C) Fluorescent images of phalloidin stained cultures at various timepoints after Moderate injury severities. (D) Total area of phalloidin-positive cytoplasm at various timepoints after Moderate injury. Groups that do not share a letter are significantly different (Kruskal-Wallis test, p< 0.05).

### Piezo1 influences calcium dynamics and redistributes after injury

The Piezo1 agonist Yoda1 increased calcium dynamics in astrocyte cultures (Fig. 4A). Group differences were assessed using a non-parametric Kruskal-Wallis test. This analysis showed a significant overall effect of treatment (p < 0.05). Post hoc pairwise comparisons were conducted using a Bonferroni correction (p<0.05). These effects were found to scale with concentration, with the most pronounced response observed at 20 µM (Fig. 4). These results show that Piezo1 is expressed and functionally active in these astrocytes.

**Figure 4:**
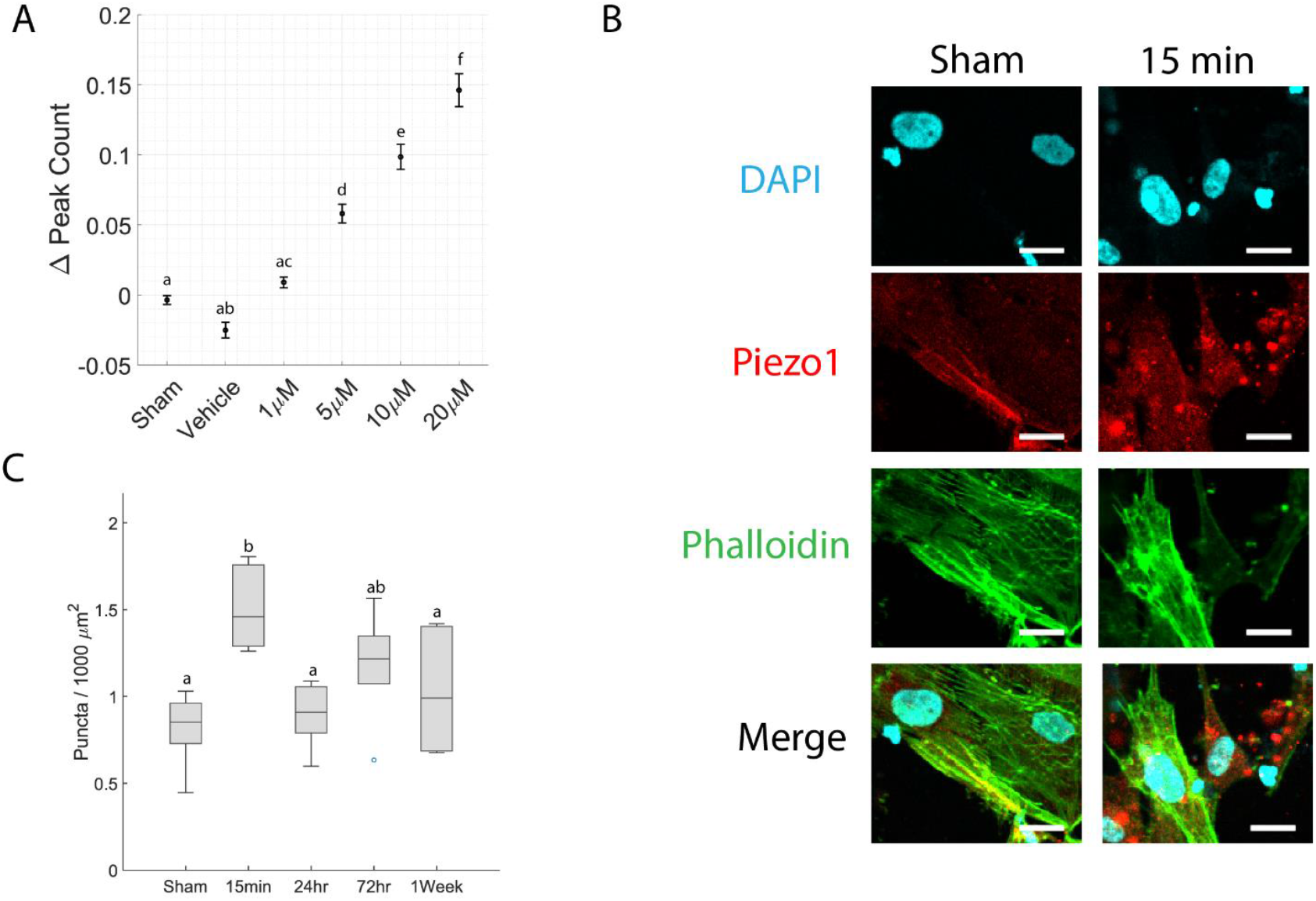
Piezo1 response to pharmacological activation and mechanical stretch. (A) Change in cell-specific calcium peak count following Yoda1-induced Piezo1 activation. Data points represent group means, with error bars indicating standard error (N=5 wells, n>=94 cells). (B) Immunostaining in sham cultures and Moderate injury cultures 15 minutes after injury (scale bar = 20μm). (C) Area density of Piezo1 puncta at various timepoints after Moderate injury. Box plots display the median and interquartile range (25th–75th percentiles).Whiskers represent the range of values within 1.5 times the interquartile range. Hollow circle represents outlier data (N=6 wells, n>=94 cells). Groups that do not share a letter are significantly different (Kruskal-Wallis test, p< 0.05).

### Trauma induces transcriptional changes in astrocytes

There were robust transcriptional alterations in response to trauma in astrocytes. Principal component analysis (PCA) shows a clear separation between the trauma and sham samples (Fig. 5A). Differential gene expression (DEG) analysis identified 196 DEGs that met the pFDR < 0.05. Of those, 184 genes were upregulated, and 12 genes were downregulated (Fig. 5B). Downregulated genes were significantly enriched for mitochondrial and oxidative metabolic processes, including oxidative phosphorylation (padj = 3.24x10^-10^), aerobic respiration (padj =3.24x10^-10^), and ATP synthesis coupled electron transport (padj =9.19x10^-9^). NHGRI-EBI GWAS-catalog enrichment analysis revealed associations with neurological traits such as Alzheimer’s disease or high density lipoprotein (HDL) levels (padj =0.012), cortical thickness (padj =0.028), bipolar I disorder (padj =0.033), and bipolar II disorder (padj =0.049) (Fig. 5C). The enrichments were tested separately for the gene ontology biological processes (GO BP) and GWAS traits, and FDR correction was applied within each gene set collection. To further evaluate trauma induced astrocyte responses, we examined expression of genes previously implicated in TBI (Fig. 5D). Trauma significantly (p<0.05) increased expression of APP, ELAVL2, and KCNJ2 and decreased expression of ELVAL1 and PIEZO1, although these did not reach FDR significance in our broader analysis. No significant changes were observed for GFAP and SLC6A11.

**Figure 5:**
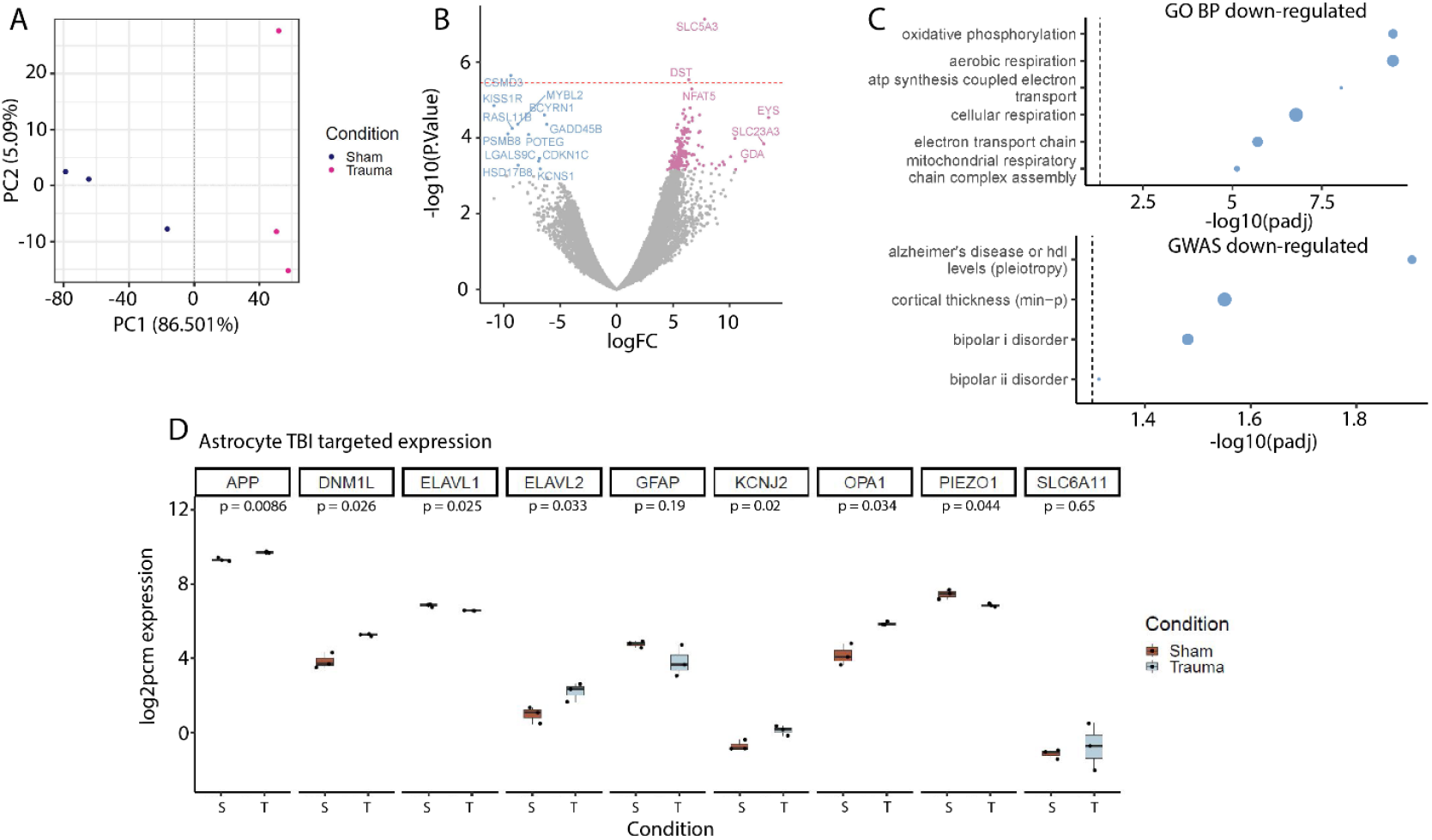
RNA-seq analysis of trauma-induced hiPSC-derived astrocytes. (A) PCA of RNA-seq separates data across sham (dark blue) and trauma (magenta) conditions. (B) Expression log fold change (difference in expression between sham and trauma conditions) was plotted against –log_10_(p-value) for each gene. Genes meeting FDR significance are color coded based on directionality: significant upregulation is pink and significant downregulation is blue. Dashed red line shows Bonferroni significance threshold (p = 3.55x10^-6^). (C) Gene ontology (GO) biological process (BP) and GWAS trait enrichments of downregulated DEGs (n=12). Terms are plotted against significance –log_10_(adjusted p-value) with dashed line representing threshold value (adjusted p-value < 0.05). Dot size reflects the proportion of genes overlapping each gene set relative to the overall number of genes in that set. (D) Log2 counts per million expression of targeted TBI genes across sham (red) and trauma (blue) conditions. P-values from two-tailed T-tests are indicated for each comparison. S and T indicate sham and trauma, respectively.

## Discussion

Mechanical trauma suppressed calcium dynamics in hiPSC-derived astrocytes. This finding is important because loss of spontaneous calcium dynamics in astrocytes can initiate neurodegenerative pathology, linking TBI to chronic neurodegeneration [44]. It is also consistent with evidence that stretch triggered calcium influx in rat astrocytes [45]. An important distinction between the cited study of rat astrocytes and this study is that rat astrocytes did not exhibit spontaneous calcium dynamics whereas hiPSC-derived astrocytes do exhibit such dynamics, both in this study and another recent publication [46]. Calcium dynamics returned to normal 7 days after injury, following the time course of mitochondrial membrane potential recovery (Fig 2B,C).

Our stretch injury model produced a repeatable injury phenotype with controllable severity in hiPSC-derived astrocytes that mimicked some phenotypes seen in prior *in vitro* studies. The first step towards achieving a repeatable injury phenotype with controllable severity was creating a repeatable strain with controllable magnitude in the membrane upon which astrocytes are cultured. Our *in vitro* stretch injury model induces pathology by pressing a cylindrical indenter into the bottom of circular PDMS membrane and forcing it to stretch equibiaxially in a horizontal plane [24], [25], [47]. This study used indentation depths of 0.5, 1.5, 2.5, and 3.2 mm and they repeatably induced peak strains of 2%, 20%, 57% 81% respectively (see Table 1). These four levels of stretch were labeled as Mild, Moderate, Severe, and Extreme. Prior studies of *in vitro* astrocyte stretch injury found a range of mild to severe pathological responses when stretching the cell culture membrane by approximately 20 to 70% [48], [49], [50]. Therefore, we are confident that the range we chose is wide enough to include a mild trauma that should not cause substantial cell death and a catastrophic trauma that should devastate the culture. 15% strain is the threshold for inducing concussion in reconstructions of impacts in American football players [51] so 20% strain is in the concussive range. Biomechanical tolerance studies estimate a 50% chance of severe TBI when at least 5% of the brain undergoes a maximum principal strain of 60.8% [52] so 57% strain is a reasonable proxy for severe injury. Observations that reproduced findings in prior *in vitro* studies of trauma in astrocytes included correlation of injury severity with cell death [48] and mitochondrial membrane depolarization [53]. In the time course experiment, the mitochondrial membrane potential increased slightly at the shortest time point studied before declining at 24 hours post-injury and then partially recovering by 72 hours post-injury (Fig. 2A,B). This finding aligns with previous reports showing that another acute stressor (oxygen-glucose deprivation (OGD)) [54], triggers an acute mitochondrial hyperpolarization followed by depolarization [55].

Yoda1, a specific agonist of Piezo1, increased calcium dynamics in astrocytes (Fig. 4A), demonstrating that Piezo1 channels were present and functionally influential in these cells. The distribution of Piezo1 became more punctate immediately after injury (Fig. 4B). Piezo1 channels have been reported to cluster in membrane invaginations [56], around focal adhesions [57], and at cell-cell junctions [58]. Our data cannot distinguish between these scenarios. However, the sudden decline in cell area after injury coincides with clustering of Piezo1 channels (Fig. 3D and Fig. 4C). Therefore, we hypothesize that membrane invaginations drive transient clustering of Piezo1 after injury. In this model of the process, injury destroys many of the focal adhesions that anchor the cell to the substrate, releasing the plasma membrane from its typical taut state, and allowing it to collapse into a more wrinkled, invaginated state with attendant clustering of Piezo1. As the cell rebuilds focal adhesions and stress fibers and restores its normal spread morphology with associated tension in the plasma membrane, these invaginations would likely be eliminated, explaining the reduction in Piezo1 puncta at the 24-hour timepoint. The functional significance of Piezo1 puncta is also unclear. Computer simulations predict that clustering would bias Piezo1 channels towards activation [59], but an experimental study concluded that Piezo1 channels did not function differently when they clustered [60]. Further experiments are needed to confirm or refute these speculations.

RNA sequencing gene ontology analysis revealed multiple processes associated with metabolism as being altered by trauma, consistent with observations in clinical and animal studies of TBI [61], [62]. Notably, our results confirm the important role that oxidative stress plays in TBI pathology [63]. The upregulation of cortical thinning pathways also agrees with clinical evidence [64]. Inspection of results for individual genes relevant to TBI pathology found statistically significant changes that agreed with prior literature. For example, amyloid precursor protein is used clinically as a biomarker of TBI [65], and the associated *APP* gene was upregulated in this model (Fig. 5D). *KCNJ2* was upregulated in a human organoid model of TBI [66] and was also upregulated in this human *in vitro* model (Fig. 5D). Perturbation of genes for RNA-binding proteins in the ELAVL family by injury is significant because these proteins influence inflammation after neurotrauma [67]. Taken together, these results indicate that many important mechanistic questions about TBI could be addressed with this hiPSC-derived astrocyte model.

Several limitations should be considered when interpreting the results of this study. The hiPSC-derived astrocytes were all derived from a single donor. Astrocytes did not interact with neurons, microglia, or other neural cell types during these experiments, precluding testing of hypotheses about how activity in astrocytes influences activity in adjacent cells and vice versa. Finally, hiPSC-derived cells most resemble fetal primary human astrocytes more than adult primary human astrocytes, which diminishes their relevance to the study of brain injuries that occur after birth. However, our team has previously shown that gene x environment stress interactions can be conserved between hiPSC-derived models and human brain [68].

In summary, mechanical stretch injury temporarily disrupts spontaneous calcium dynamics, reduces cell area, and redistributes Piezo1 channels in hiPSC-derived astrocytes, establishing them as a platform for the study of post-TBI morbidity.

## Supplementary data

**Figure S1:**
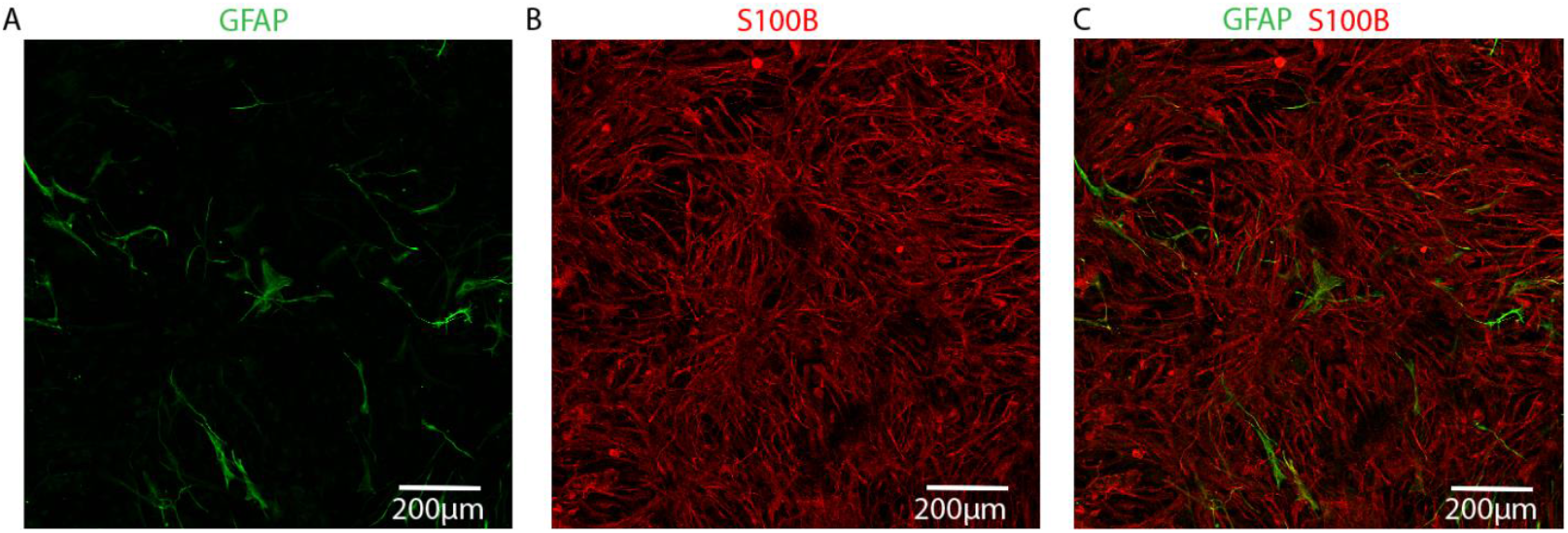
Immunostaining for (A) GFAP (B) S100B and (C) Merged in 21 day old astrocytes-Images are representative images of 3 wells.

**Figure S2:**
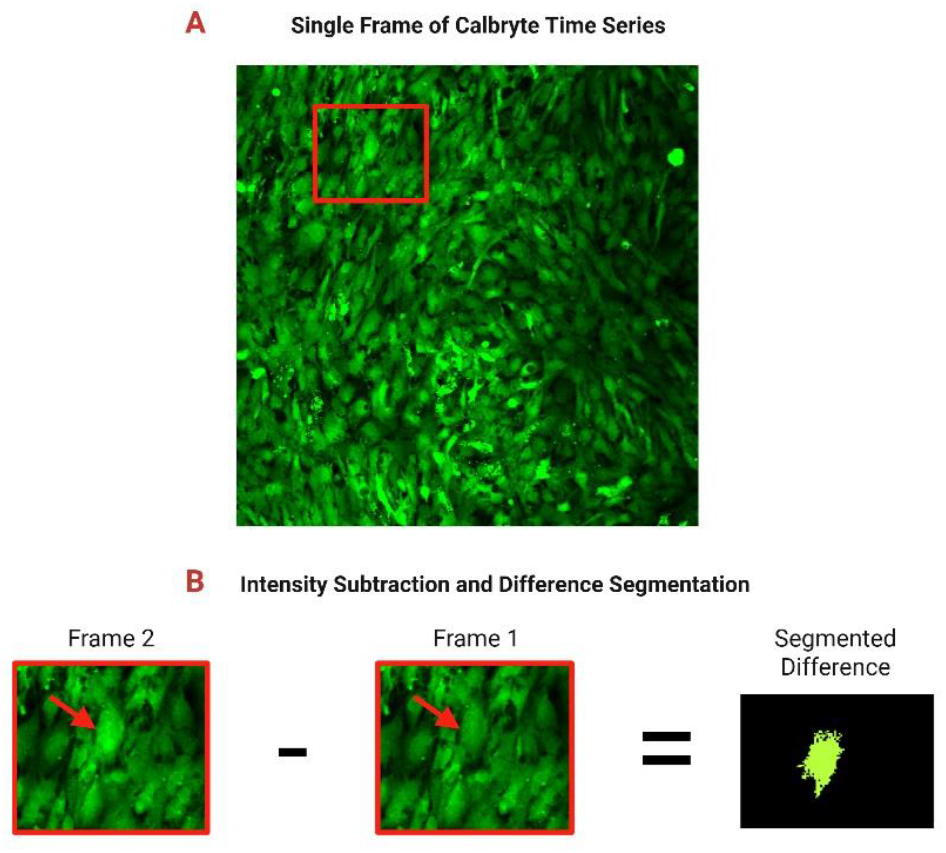
Workflow of asynchronous activity index (AAI) segmentation (**A**) Single frame from a Calbryte time series of a sham culture. Red box indicates the region selected for visualization (B) Intensity subtraction and difference segmentation. The cropped region from Panel A (left) and the corresponding region from the previous frame (middle) are shown with red arrows indicating areas of fluorescence change. Regions with larger intensity differences are segmented into discrete objects (right) for further analysis.

**Fig S3.**
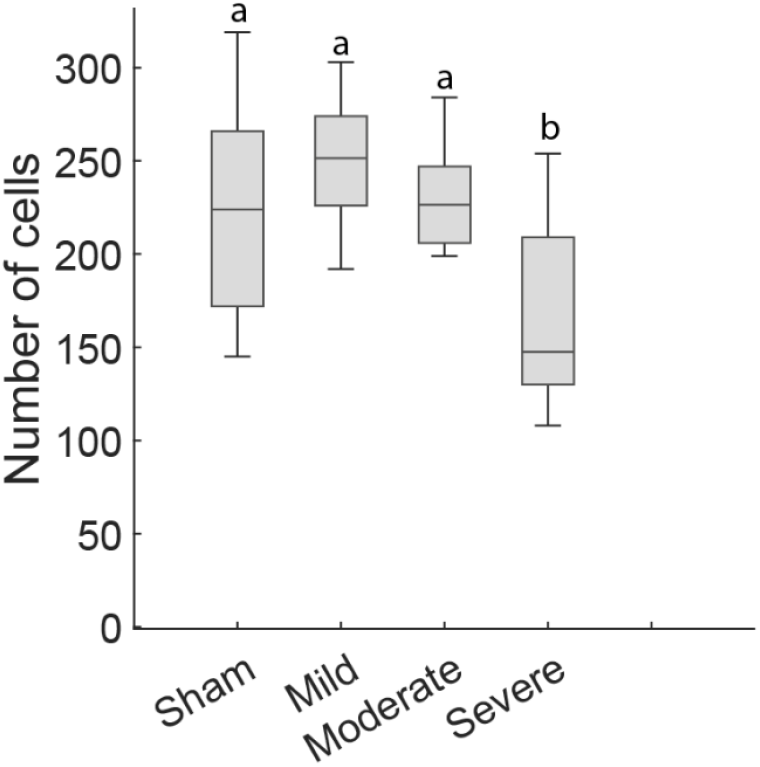
Number of cells 24 hours after injury. Box plots display the median and interquartile range (25th–75th percentiles).Whiskers represent the range of values within 1.5 times the interquartile range (N=6 wells). Groups that do not share a letter are significantly different (Kruskal-Wallis test, p< 0.05).

**Fig S4.**
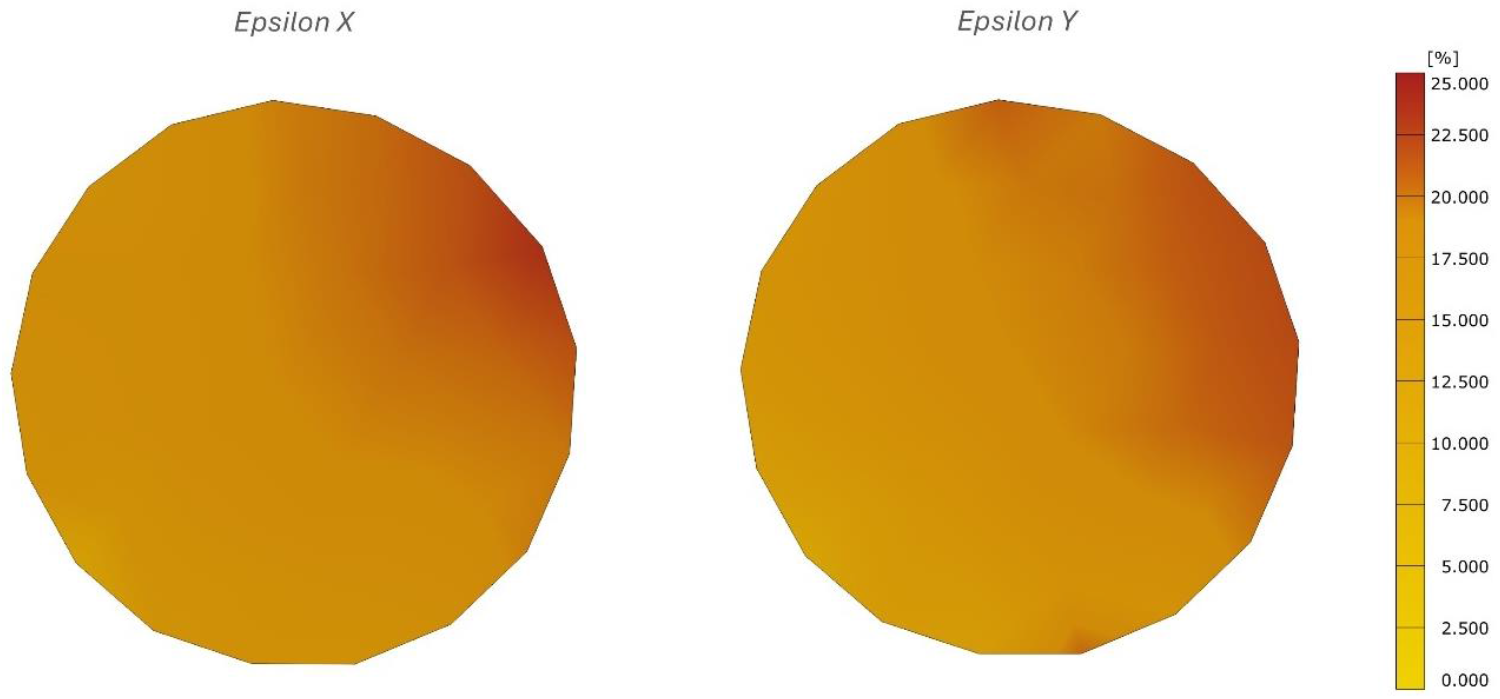
Peak strain distributions in X and Y directions for a 1.5 mm deep indentation. (Representative image of 21 wells).

